# Engineering precise adenine base editor with infinitesimal rates of bystander mutations and off-target editing

**DOI:** 10.1101/2022.08.12.503700

**Authors:** Liang Chen, Shun Zhang, Niannian Xue, Mengjia Hong, Xiaohui Zhang, Dan Zhang, Jing Yang, Sijia Bai, Yifan Huang, Haowei Meng, Hao Wu, Changming Luan, Biyun Zhu, Gaomeng Ru, Meizhen Liu, Mingyao Liu, Yiyun Cheng, Chengqi Yi, Gaojie Song, Liren Wang, Dali Li

## Abstract

Adenine base editors (ABEs) catalyze A-to-G transitions showing broad applications, but their bystander mutations and off-target editing effects raise the concerns of safety issues. Through structure-guided engineering, we found ABE8e with an N108Q mutation reduced both adenine and cytosine bystander editing, and introduction of an additional L145T mutation (ABE9), further refined the editing window to 1-2nt with eliminated cytosine editing. Importantly, ABE9 induced very minimal RNA and undetectable Cas9-independent DNA off-target effects, which mainly installed desired single A-to-G conversion in mouse and rat embryos to efficiently generate disease models. Moreover, ABE9 accurately edited A_5_ position of the protospacer sequence in pathogenic homopolymeric adenosine sites (up to 342.5-fold precision than ABE8e) and was further confirmed through a library of guide RNA-target sequence pairs. Due to the minimized editing window, ABE9 could further broaden the targeting scope for precise correction of pathogenic SNVs when fused to Cas9 variants with expanded PAM compatibility.

## Introduction

DNA base editors are innovative genome editing tools catalyzing efficient base conversions without creating DNA double strand breaks (DSBs) or a requirement for donor DNA templates^1^. Cytosine base editors (CBEs) are composed of Cas9 nickase (nCas9) and a cytosine deaminase domain to catalyze specific C•G-to-T•A transitions with the presence of a uracil glycosylase inhibitor (UGI)^2^. Similarly, adenine base editors (ABEs) were developed by fusion of nCas9 to a wild-type or an evolved TadA (eTadA) (originally a transfer RNA (tRNA) adenine deaminase in *Escherichia coli*) to efficiently generate A•T-to-G•C conversions^3^. Unlike CBEs which also induce C to non-T side-products and indels due to activation of the base excision repair pathway, the first generation of ABEs (like ABE7.10) produces pure A-to-G conversions without inducing significant indels (typically ≤0.1%)^3^. More importantly, ABE7.10 rarely induces Cas9-independent off-target DNA editing which has been reported in CBE-treated cells and embryos^4, 5^. These excellent features make ABEs promising tools for future clinical applications.

Great efforts have been made to improve the performance of ABEs. As TadA is a tRNA adenine deaminase, numerous occurrences of random RNA off-target editing have been reported^6-8^. Through the introduction of point mutations in wtTadA/eTadA or using only an engineered eTadA, several versions of ABEs, such as ABEmax-F148A^7^ (an F148A mutation introduced to both TadA and eTadA), ABEmax-AW^8^ (with TadA E59A and eTadA V106W mutations) and SECURE-ABEs^6^ (with eTadA K20A/R21A or V82G mutations) exhibited minimized off-target edits. To improve the editing efficiency and targeting scope, two new groups of ABE variants, ABE8e^9^ and ABE8s^10^, have been developed through molecular evolution of the eTadA monomer. ABE8e is the most efficient and compatible ABE variant whose activity exhibits a 3- to 11-fold improvement compared with ABE7.10, while it also expands the editing window^9^. ABE8e and ABE8s also showed quite high editing efficiencies in the livers of mice and non-human primates^11^ or hemopoietic stem cells from sickle cell anemia patients^12^, demonstrating their glorious potential for gene therapeutics. However, with the increase of deamination activity, ABE8e exhibits significant Cas9-independent DNA and RNA off-target editing^9, 13, 14^.

Although ABE8 variants are highly efficient, the editing window is also expanded with significant editing rates on the bystander adenines^9, 10, 15^. Moreover, several studies have shown that ABE7.10 exhibits cytosine deamination activity which enables C-to- T/G/A conversions with a preference for TCN motif, demonstrating that ABEs also induce undesired bystander cytosine mutations in cell lines and animal embryos^16-18^. It is critical to eliminate both adenine and cytosine bystander effects and Cas9-independent off-targeting editing of ABEs, especially for clinical applications. In this study, through structure-based engineering, we generated ABE9 which accurately catalyzed A-to-G conversions within a 1-2nt editing window without inducing C-to-T conversions in cells and rodent embryos. We also demonstrated it precisely corrected pathogenic SNVs, especially in homopolymeric adenosine sites with infinitesimally small rates of Cas9-independent RNA and DNA off-target effects.

## Results

### Structure-based molecular evolution of TadA-8e

ABE8e, whose deaminase component is a multiple-turnover enzyme with high processivity^19^, edits more positions than previously reported ABEs^9^. We also confirmed that adenines in positions 3-12 were efficiently edited by ABE8e, suggesting a much wider editing window than ABEmax (Extended Data Fig. 1a). ABE8e also exhibited elevated cytosine bystander editing effects and increased Cas9-independent DNA off-target editing through a more sensitive orthogonal R-loop assay which uses SaCas9 nickase instead of dSaCas9^9, 20, 21^ (Extended Data Fig. 1b,c). These elevated rates of undesired ABE8e editing effects encourage us to further optimize it for more accurate editing.

To increase its accuracy, we intended to evolve the TadA-8e based on its DNA binding cryo-electron microscopy structure^19^ (PDB: 6VPC). The structure suggests that three nucleotides including the editing base (labelled “0” in Fig. 1a) and the ones before and after it (labelled “-1” and “+1”, respectively) of the substrate are important for recognition by the deaminase. We hypothesized that mutating these residues that interacted with either the bases or the backbone of the substrate would change the environment of the binding pocket as well as the accessibility to the substrate. It might eventually reduce the non-specific binding and narrow down the editing window. Moreover, according to the apparently different size and electrophilicity of the purine ring (A) compared to the pyrimidine ring (C), these mutations would change the substrate selectivity of TadA-8e deaminase. Residues included the E27-V28-P29 loop and F148 that inserted into a valley formed by the “0” and “+1” bases; the F84, N108, L145, and Y149 that inserted into the other valley formed by the “0” and “-1” bases, and the P86/H57 that was adjacent to the editing base (Fig. 1a). Thus, 10 residues were individually mutated to remove the large side chain (*e*.*g*., F84T, F148A) or add a bulky residue (*e*.*g*., V28F, P29W), or change between non-polar and polar residues (*e*.*g*., L145T, V28N, N108V). Some highly conserved positions adjacent to the pocket (*e*.*g*., E27-P29, L145, F148) were also included to maximize the possibility of developing a precise editing tool.

**Fig. 1.**
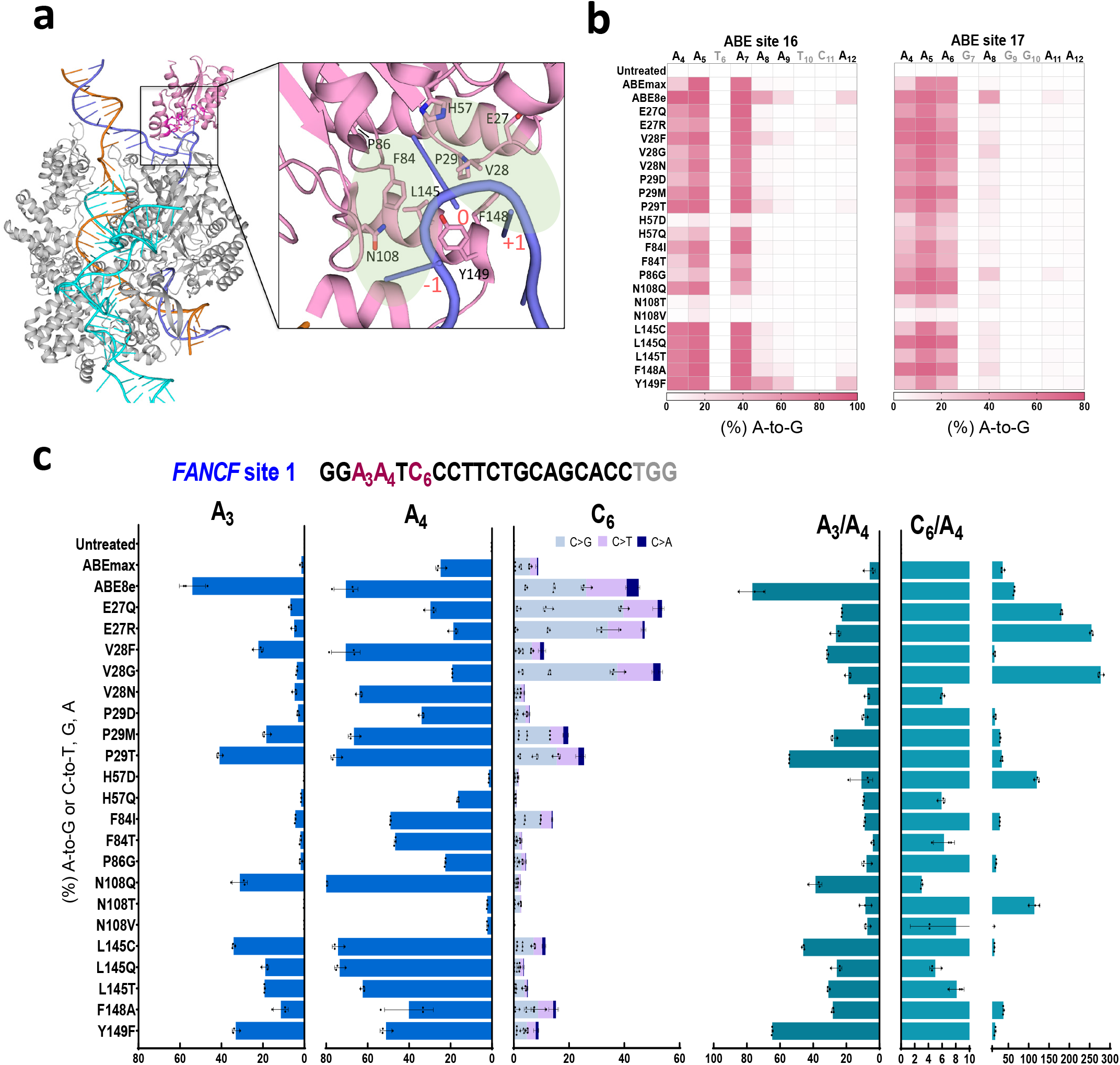
Structure-based molecular evolution of TadA-8e. **a**, The schematic diagram of the interplays of TadA-8e (pink) with the ssDNA substrate (light blue sticks) (PDB: 6VPC). Complementary strand DNA is in orange, non-complementary strand DNA is in light blue, Cas9n is in gray, and sgRNA is in cyan. Amino acids reacting with the substrate DNA are labeled on the enlarged image. **b**, The A-to-G base editing efficiency of ABE8e or ABE8e variants at 2 endogenous genomic loci containing multiple adenosines (ABE site 16 and ABE site 17) in HEK293T cells. The heatmap represents an average editing percentage derived from three independent experiments with editing efficiency determined by HTS. **c**, Base editing efficiency of ABE8e or ABE8e variants at an endogenous genomic locus (*FANCF* site 1) for both adenine and cytosine editing in HEK293T cells. A_3_/A_4_ means the ratio of A_3_ editing to A_4_ editing, and C_6_/A_4_ means the ratio of C_6_ editing to A_4_ editing. Data are mean ± s.d. (n = 3 independent experiments). Statistical source data are provided online.

Following the above principles, 21 point mutations were constructed in TadA-8e and the activity was determined on three target sites. Deep-sequencing data of the first two targets with multiple adenines showed that the majority of the mutations reduced the editing window with a comparable or slightly decreased A-to-G efficiency compared to ABE8e, while H57D, H57Q, N108T and N108V mutations dramatically reduced the activity. In contrast to ABEmax^7^, the introduction of an F148A mutation in TadA-8e did not narrow the editing window (Fig. 1b). On the third target site previously used for evaluation of cytosine bystander mutations^13, 16^, ABEmax and ABE8e induced lots of cytosine mutations (8.83% and 45.20% in average) while V28F, V28N, N108Q, L145C, L145T and L145Q mutations exhibited high A-to-G activity on A_4_ with greatly reduced cytosine conversions (ranging from 2.43% to 11.47%) (Fig. 1c). After comparing with the editing efficiency ratio of A_3_/A_4_ or C_6_/A_4_, we found that ABE8e-N108Q had the best performance and was selected for further investigations (Fig. 1c).

### ABE8e-N108Q reduces bystander adenine and cytosine editing

To further profile the performance of ABE8e-N108Q, 21 endogenous targets were tested in HEK293T cells by high-throughput sequencing (HTS). The first batch of 12 target sites contained multiple adenosines and the other 9 targets contained mixed adenines and cytosines in the editing window. ABE8e was highly efficient (>50%) between positions A_3_ to A_8_ and considerable editing was also observed in a very lateral position such as A_2_ or A_13_, but ABE8e-N108Q mainly edited A_4_-A_7_ with almost no editing on A_9_ to PAM-proximal positions (Fig. 2a and Extended Data Fig. 2a). In the remaining 9 targets, we found that in addition to the TCN motif, ABE8e also edited cytosines in CCN, GCN and ACN motifs (Fig. 2b). ABE8e induced cytosine base conversions up to 39.23% (*SSH2*-sg10), with the highest efficiency on C_6_ with an average rate of 18.02% (Fig. 2b,c and Extended Data Fig. 2b). In contrast, a significant decrease of cytosine conversions was observed in ABE8e-N108Q-treated cells with an average editing efficiency of 5.79% on C_6_, although its cytosine deaminase activity was not fully eliminated (Fig. 2c). Based on all 21 target results, ABE8e-N108Q exhibited an identical A-to-G efficiency with ABE8e at the highest positions (82.1% vs 82.74% on A_5_ and 83.62% vs 83.13% on A_6_), but the major editing window was reduced to A_4_-A_7_ (Fig. 2d). Similar to ABE8e, ABE8e-N108Q minimally induced indels on the selected target sites (Fig. 2e). Together, ABE8e-N108Q is highly efficient with significantly reduced adenine and cytosine bystander mutation effects.

**Fig. 2.**
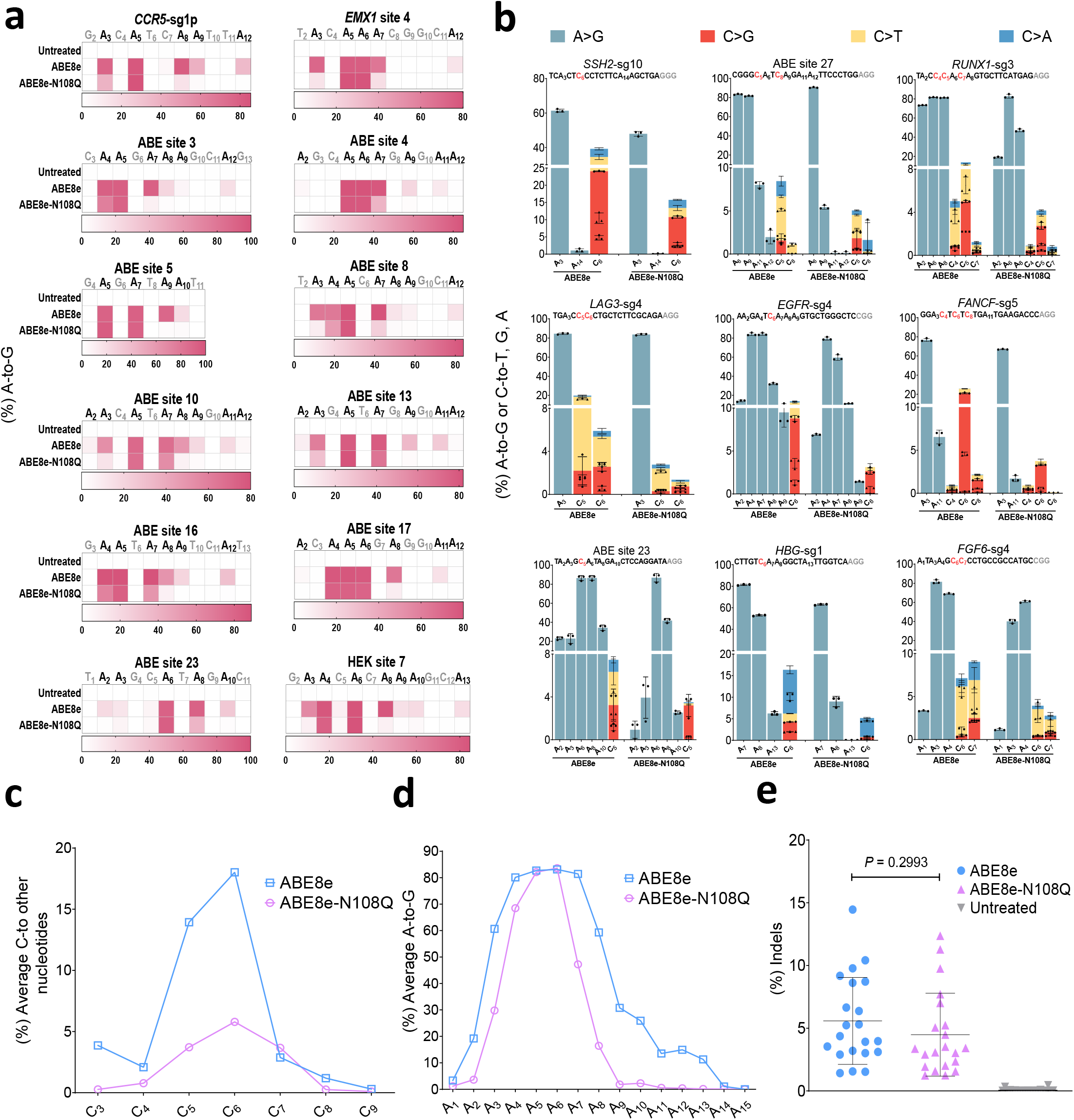
Characteristics of ABE8e-N108Q in HEK293T cells. **a**, The editing efficiency of ABE8e or ABE8e-N108Q was examined at 12 endogenous genomic loci containing multiple As in HEK293T cells. The heatmap represents the average editing percentage derived from three independent experiments. **b**, The editing efficiency of ABE8e or ABE8e-N108Q was examined at 9 endogenous genomic loci containing an NCN motif in HEK293T cells. Data are mean ± s.d. (n = 3 independent experiments). **c**, Average C-to-T/G/A editing efficiency of ABE8e or ABE8e-N108Q at the 9 target sites in **b. d**, Average A-to-G editing efficiency of ABE8e or ABE8e-N108Q at the 21 target sites in **a**,**b. e**, Frequency of indel formation by ABE8e or ABE8e-N108Q at the 21 target sites in **a**,**b**. Each data point represents the average indel frequency at each target site calculated from 3 independent experiments. Error bars and *P* value are derived from these 21 data points. *P* value was determined by two-tailed Student’s t test. In **c** and **d**, data represent averages from three independent experiments. Statistical source data are provided online.

### Single adenine to guanine transition by further evolution of ABE8e-N108Q

Although ABE8e-N108Q exhibits a smaller editing window and fewer cytosine edits, we pursued a more accurate ABE featuring a single nucleotide window and complete elimination of cytosine editing activity. We assumed that introducing more mutations in ABE8e-N108Q would further reduce bystander editing. As shown in Fig. 3a, the combination of N108Q with an additional single mutation on residues E27, P29, F84 or L145 exhibited a very stringent editing window even to a single adenine at the A_5_ position. Among the variants, ABE8e-N108Q/L145T showed single nucleotide editing window and high activity at these two sites, and we named it ABE9 as one more mutation was introduced into ABE8e. As tested through 12 endogenous sites, we found that ABE9 was most efficient at A_5_ with comparable activity to ABE8e and ABE8e-N108Q on most of the tested targets (Fig. 3b). Importantly, its activity at the adjacent A_4_ or A_7_ position was dramatically reduced or even eliminated at 8 of these 12 targets, and single adenine editing was observed at half of tested sites (Fig. 3b). Using the editing rate of the most efficient position to divide by the second highest position, we further confirmed that ABE9 was the most accurate variant which showed up to 12.7-fold (5.1-fold in average) discrimination of the two most efficient adenine positions than ABE8e-N108Q (Fig. 3c). Collectively, we developed a more accurate ABE variant ABE9, which showed a stringent and steep editing window of 1-2 nucleotides at A_5_ or A_6_ (Fig. 3d and Extended Data Fig. 3a). As expected, the indel rates of ABE9 are comparable or even slightly reduced than ABE8e and ABE8e-N108Q (Extended Data Fig. 3b).

**Fig. 3.**
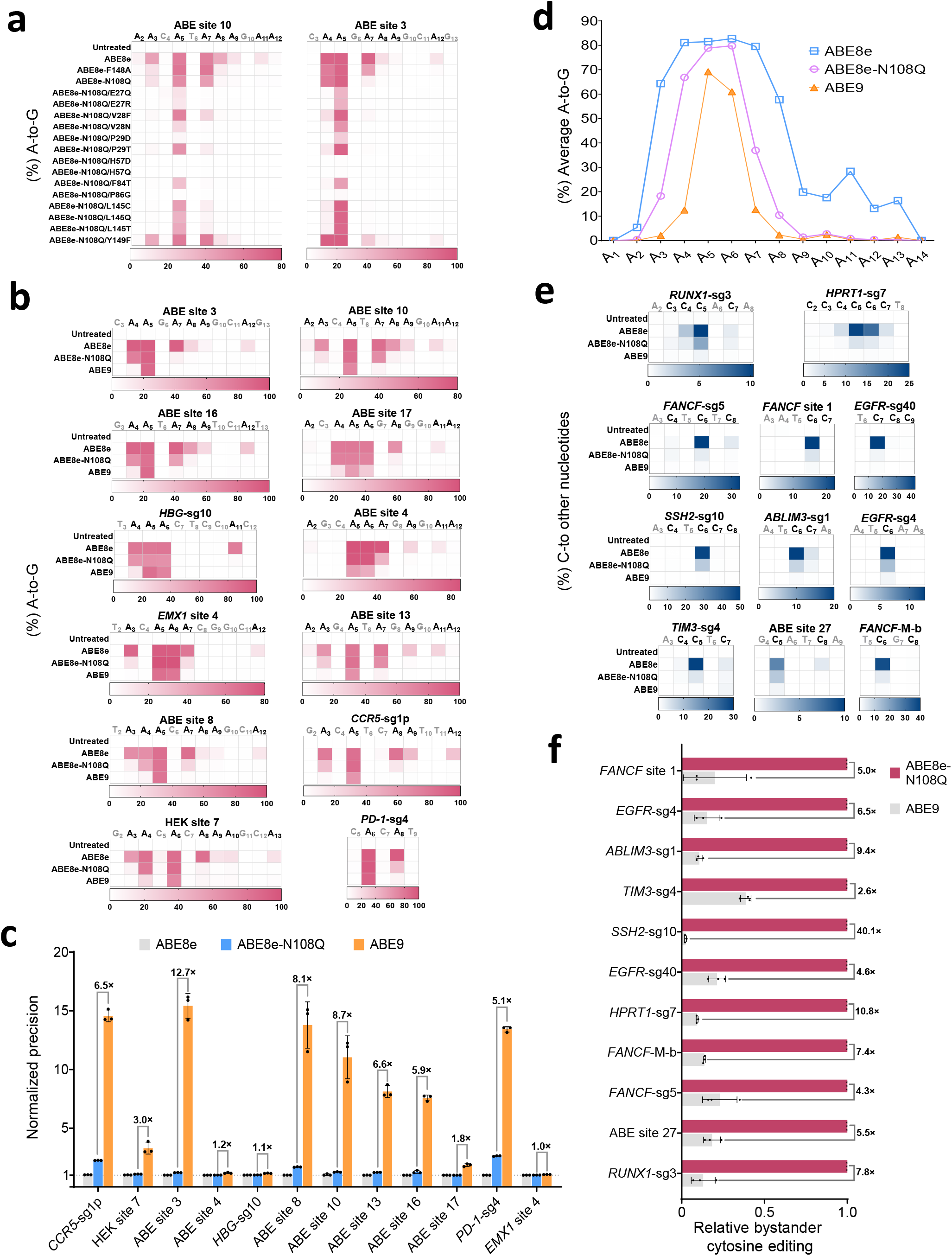
Evolution and characterization of single A-to-G base editor. **a**, The A-to-G base editing efficiency of ABE8e-N108Q and its combination variants at 2 endogenous genomic loci containing multiple As (ABE site 10 and ABE site 3) in HEK293T cells. **b**, The A-to-G editing efficiency of ABE8e-N108Q or ABE9 was examined at 12 endogenous genomic loci containing multiple As in HEK293T cells. **c**, The normalized precision (ABE8e is used for standardization) is defined as the highest / second-highest A-to-G base editing of ABE8e-N108Q or ABE9 at the 12 target sites in **b**. Data represent mean ± s.d. from three independent experiments. **d**, Average A-to-G editing efficiency of ABE8e-N108Q or ABE9 at the 12 target sites in **b**. Data represent mean from three independent experiments. **e**, The C-to-T/G/A editing efficiency of ABE9 was examined at 11 endogenous genomic loci containing multiple Cs in HEK293T cells. **f**, The normalized ratio (ABE8e-N108Q is used for standardization) of the highest C-to-T/G/A editing efficiency of ABE8e-N108Q or ABE9 at 11 target sites in **e**. Data represent mean ± s.d. from three independent experiments. In **a, b** and **e**, the heatmap represents average editing percentage derived from two or three independent experiments. Statistical source data are provided online.

Next, 11 target sites were employed to evaluate the cytosine bystander mutation rate. As shown in Fig. 3e, ABE9 did not edit Cs in 10 of these 11 targets whereas ABE8e and ABE8e-N108Q induced considerable edits on all targets tested (efficiency lower than 1% considered as no editing). The highest cytosine conversion activity of ABE9 detected was 1.5% on C_7_ of the *TIM3*-sg4 target but ABE8e and ABE8e-N108Q catalyzed 29.6% and 3.9% conversions on C_5_, respectively. According to statistical analysis of the cytosine editing rate of the most efficient position, ABE9 strikingly decreased the cytosine bystander mutation rate by 2.6- to 40.1-fold (9.5-fold on average) normalized to that of ABE8e-N108Q (Fig. 3f).

### Off-target analysis of ABEs in mammalian cells

To evaluate Cas9-dependent off-target activity, 44 potential off-target sites from 5 sgRNA targets were analyzed, including 17 known off-target sites identified by GUIDE-seq or ChIP-seq^3, 22^ and 27 in silico-predicted off-target sites by Cas-OFFinder^23^. We found that ABE8e induced mild off-target editing (1.04-12.29%, 4.15% on average) at 11 sites on HEK site 2, HEK site 3 and *PD-1*-sg4 loci, while ABE9 only edited two sites with comparable on-target activity (Fig. 4a and Extended Data Fig. 4). The Cas9-independent DNA and RNA off-target editing caused by the deaminase were more unpredictable and intractable. Through an enhanced orthogonal R-loop assay^21, 22^, Cas9-independent DNA and RNA off-target effects of ABE9 were greatly reduced compared to ABE8e (Fig. 4b,c and Extended Data Fig. 5). Amazingly, the off-target activity of ABE9 was lowered to near-background levels (mean <0.3%) (Fig. 4b), indicating that it eliminated unpredictable DNA off-target activity. Through whole genome mRNA profiling analysis, we found that RNA off-target effects of ABE9 were reduced to background level and displayed 726.1- and 117.1-fold reduction compared to ABE8e and ABE8e-N108Q, respectively (Fig. 4c). These results demonstrate that ABE9 is highly specific with infinitesimal rates of unpredictable DNA and RNA off-target activity.

**Fig. 4.**
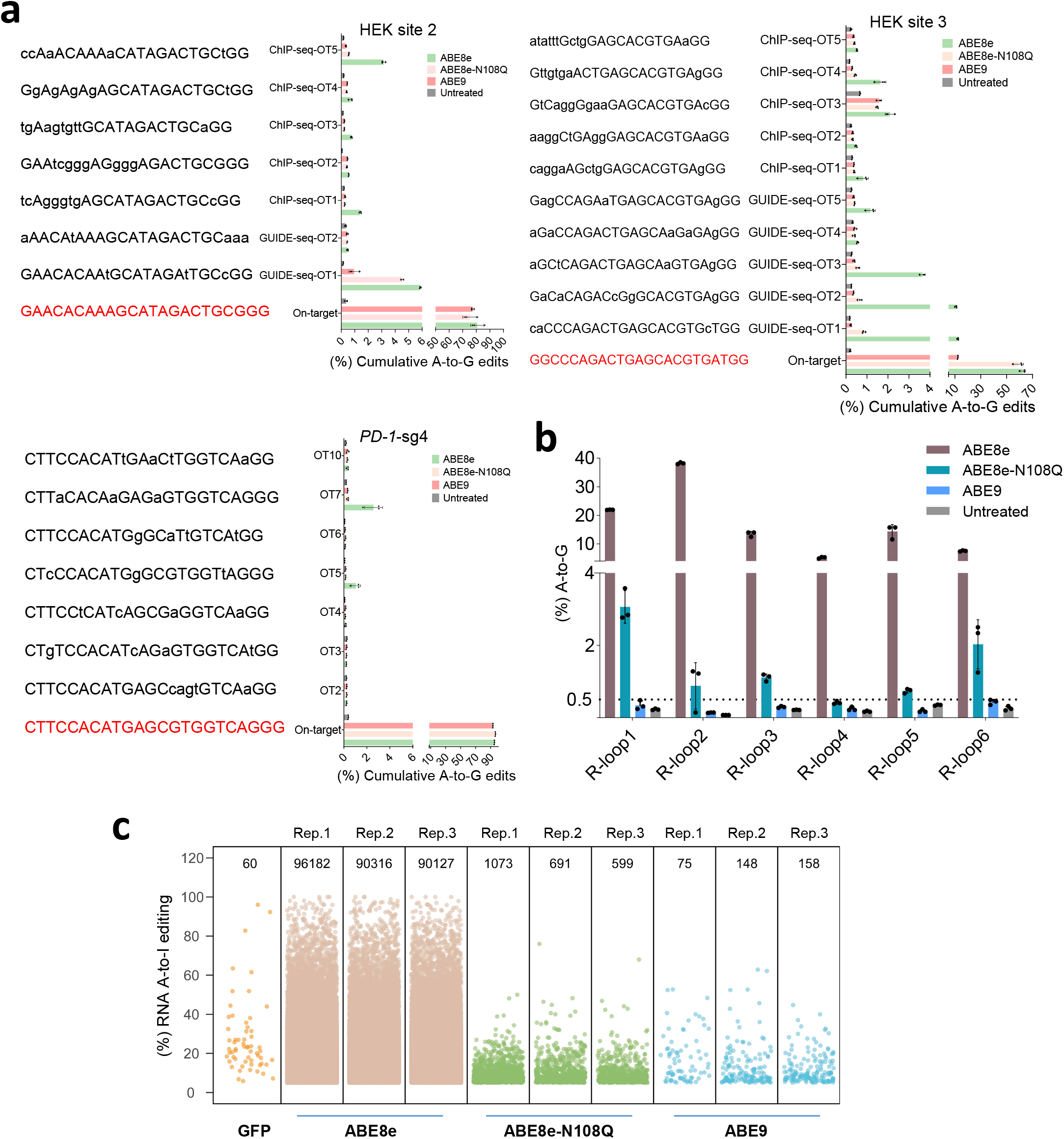
Off-target mutation assessment of ABE9. **a**, Cas9-dependent DNA on and off-target analysis of the indicated targets (HEK site 2, HEK site 3 and *PD-1*-sg4) by ABE8e, ABE8e-N108Q and ABE9 in HEK293T cells. **b**, Cas9-independent DNA off-target analysis of ABE8e, ABE8e-N108Q and ABE9 using the modified orthogonal R-loop assay at each R-loop site with nSaCas9-sgRNA plasmid. **c**, RNA off-target editing activity by ABE8e, ABE8e-N108Q and ABE9 using RNA-seq. Jitter plots from RNA-seq experiments in HEK293T cells showing efficiencies of A to I conversions (y-axis) with ABE8e, ABE8e-N108Q and ABE9 or a GFP control. Total numbers of modified bases are listed on the top. Each biological replicate is listed on the bottom. In **a** and **b**, data are mean ± s.d. (n = 2 or 3 independent experiments). Statistical source data are provided online.

### Highly efficient and accurate editing by ABE9 in rodent embryos

Accurate base conversion is critical for modeling pathogenic SNVs, but ABEs or CBEs usually induced severe bystander mutations at the target sites in cells and embryos^24-26^. To test whether ABE9 could generate precise single nucleotide conversion in embryos, ABE8e or ABE9 mRNA was co-injected with sgRNA targeting the splicing acceptor site of *Tyrosinase* gene intron 3 into mouse zygotes to model albinism. Once the splicing site was destroyed (A_5_ position), exon skipping might occur to disrupt tyrosinase coding and lead to an albino phenotype (Fig. 5a). After deep sequencing of genomic DNA from F0 pups, all of the ABE8e or ABE9 injected mice contained A_5_ editing (Fig. 5b and Extended Data Fig. 6a). Notably, ABE9 selectively edited A_5_ in 88% (14 out of 16) of the pups and the other two pups bore very low (8.13% and 9.75%) simultaneous A8 conversions, but only 5% (1 out of 19) of the pups generated by ABE8e injection bearing desired A_5_ transition (Extended Data Fig. 6a,b). After analysis of total NGS reads from all F0 pups in the same group, ABE9 generated the desired A_5_ transition in 54.32% of the reads, but only 5.1% of the reads induced by ABE8e was the desired mutation (Fig. 5c). The albino phenotype in the eyes and fur color of the founders suggested that tyrosinase activity was disrupted by ABE9-induced A_5_ conversion (Fig. 5d).

**Fig. 5.**
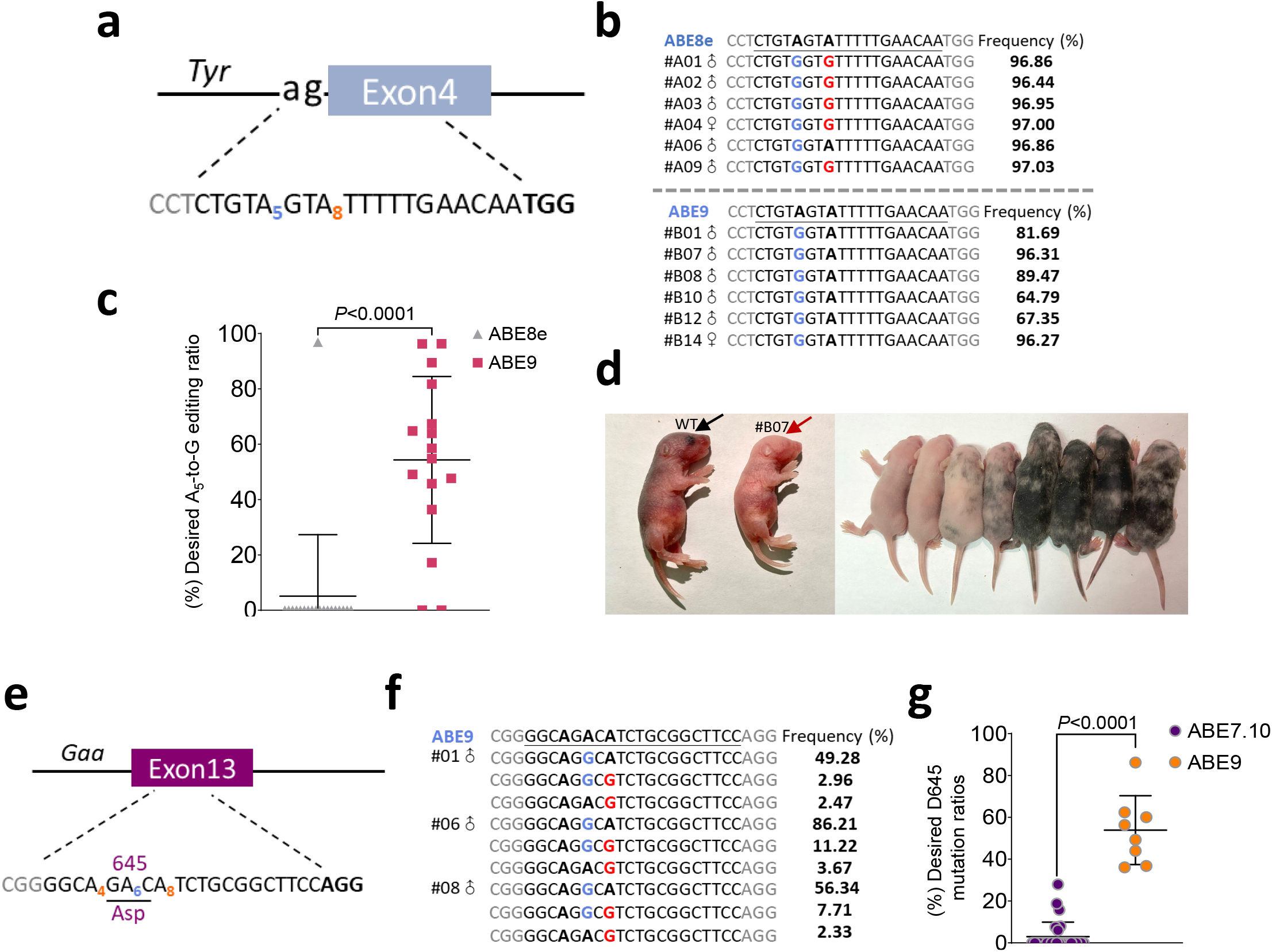
Examination of precision in rodent embryos with ABE9. **a**, The target sequence of the splicing acceptor site in intron 3 of the mouse *Tyr* gene. The “ag” sequence of the splice acceptor site is shown in black. The PAM sequence and the sgRNA target sequence are both shown in black and the PAM sequence is in bold. Target A_5_ is shown in blue with bystander A_8_ marked in red. **b**, Genotyping of representative F0 generation pups from mouse embryos microinjected with ABE8e or ABE9. The guanines converted from editable As indicate desired editing in A_5_ (blue) or undesired editing in A_8_ (red). **c**, Single A_5_-to-G conversion ratio in F0 mice induced by ABE8e or ABE9. **d**, Phenotype of F0 generated by microinjection of sgRNA and corresponding ABEs. The photo on the left was taken when the mouse was at 7 days old, while the right one was at 14 days old. WT, wildtype. **e**, The target sequence of exon 13 (dark purple) in the rat *Gaa* gene. The sgRNA target sequence where target A_6_ is shown in blue with bystander A_4_ and A_8_ marked in red is shown in black (PAM in bold). The triplet codon of D645 is underlined. **f**, Genotyping of representative F0 generation pups from rat embryos microinjected with ABE9 (desired editing in blue or undesired editing in red). **g**, Desired D645 mutation ratios in F0 rats induced by ABE7.10 or ABE9. In **b** and **f**, the percentage on the right represents the frequency of the indicated mutant alleles determined by HTS. In **c** and **g**, *P* value was determined by two-tailed Student’s t test. Statistical source data are provided online.

We further inspected the efficiency and accuracy of ABE9 in rat embryos through targeting a site with three adenines in an A_4_-A_8_ canonical editing window (Fig. 5e). As our previous data showed that only the A_6_-to-G conversion, which caused D645 mutation in *Gaa* gene identified in early-onset Pompe (glycogen storage disease type II) disease patients, leading to obvious phenotype in rats^26^. Through reanalysis of our published data, it showed that ABE7.10 only induced 6 of 28 (21%) pups bearing desired D645G mutation with the efficiency ranging from 6.04% to 27.94 % (Extended Data Fig. 6c). In contrast, ABE9 induced desired A_6_ substitution in all 8 (100%) pups with the efficiency ranging from 36.08% to 86.21% (Fig. 5f,g and Extended Data Fig. 6d). From HTS results of all 28 F0 rats treated with ABE7.10, the proportion of desired reads was only about 2.76% of all cumulative HTS reads, while ABE9 induced a 19.5-fold increase (53.77%) compared to that of ABE7.10-treated rats (Extended Data Fig. 6e), suggesting ABE9 was more efficient and accurate than ABE7.10. These data demonstrate that ABE9 is very efficient to generate highly accurate base installation in mouse and rat embryos.

### Precise correction of pathogenic mutations by ABE9

ABE generates A-to-G conversions and potentially corrects approximately half of known pathogenic SNVs in the ClinVar database, irrespective of bystander mutations^27^. To investigate the therapeutic potential of ABE9 for treating genetic diseases, 4 pathogenic SNPs with at least 4 consecutive adenines within positions 4-8 were tested, including missense mutations in *COL1A2* gene (causing autosomal-dominant osteogenesis imperfecta)^28, 29^, *CARD14* gene (causing psoriasis)^30^, *BVES* gene (causing muscular dystrophy)_31_ and *KCNA5* gene (causing common cardiac rhythm disorder)^32^. ABEs were transfected into 4 stable cell lines containing the pathogenic variants described above. For the *COL1A2* locus, ABE8e or ABE8e-N108Q did not generate considerable conversions selectively on A_5_, while ABE9 induced 34.25% desired single A-to-G conversion which was 342.5- and 21-fold higher than ABE8e and ABE8e-N108Q, respectively. Similarly, for the other three loci, ABE8e and ABE8e-N108Q only generated desired edits with frequencies of up to 2.06% and average 5.3% (0.3-11.6%), respectively, while ABE9 generated precisely corrected alleles in all four targets with an efficiency ranging from 15.53-37.22% (mean 30.19%), suggesting it was very accurate to generate single nucleotide transition (Fig. 6a and Extended Data Fig. 7a-d). These data demonstrate that ABE9 is a tremendously precise and efficient editor with the ability to correct genetic variants even in promiscuous homopolymeric sites.

**Fig. 6.**
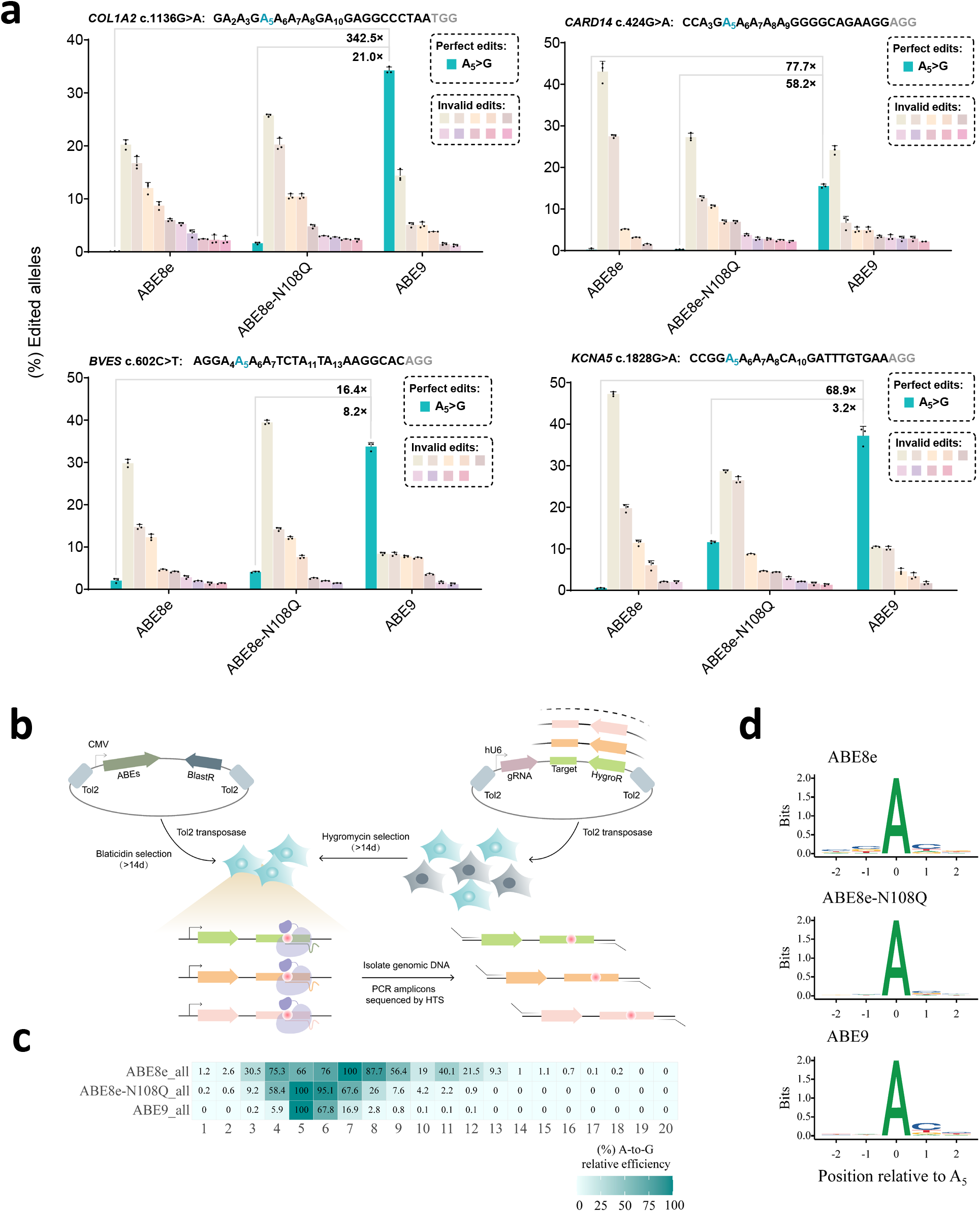
Correction of human pathogenic mutations in mammalian cells and target library analysis to unbiasedly characterize ABE9. **a**, Comparison of base correcting pathogenic mutations induced by ABEs in four stable HEK293T cell lines (*COL1A2* c.1136G>A, *KCNA5* c.1828G>A, *BVES* c.602C>T, *CARD14* c.424G>A). Base editing efficiency was determined by HTS. Data are mean ± s.d. (n = 2 or 3 independent experiments). Desired A_5_-to-G percentiles of alleles (green bar) are exhibited, while percentiles of the top ten invalid allele types are presented and percentiles of invalid allele types less than 1% are omitted. The numbers above green bars display the fold changes of ABE9 in desired A_5_-to-G percentiles compared with ABE8e and ABE8e-N108Q. **b**, Schematic of target library analysis. **c**, Analysis of relative editing efficiency of ABE8e, ABE8e-N108Q and ABE9. The heatmap represents editing efficiency computed relatively to the highest A-to-G base editing of the protospacer. Positions of the protospacer are shown at the bottom row, counting the protospacer adjacent motif (PAM) as positions 21-23. **d**, Motif visualization of ABE8e, ABE8e-N108Q and ABE9 in fifth adenine-containing cassettes. Statistical source data are provided online.

### Target library analysis of ABE9

To unbiasedly characterize the performance of ABE9, we adapted the guide-RNA target pair strategy^33, 34^ and synthesized a library of 9120 oligos with all possible 6-mers containing at least an adenine and a cytosine across positions 4 to 9 of a protospacer (see Methods). The oligo library was stably integrated into the genome of HEK293T cells via Tol2 transposon followed by stable transfection of a given base editor (Fig. 6b). We maintained an average 99% coverage of >300× per guide-target pair throughout the culturing process (Supplementary Table 4). Subsequently, the target region was amplified and sequenced at an average depth of 860 per target. The editing efficiency of the highest position in each target was considered as 100%, and the relative activity of other positions was determined comparing with the highest position. Analysis of the editing outcomes from three distinct base editors showed that ABE8e (evaluated 9059 sgRNAs) had a wide editing window ranging from positions 3-12 with a major window (>50%) from 4-9, while ABE8e-N108Q (evaluated 9071 sgRNAs) narrowed the window to positions 4-8 (Fig. 6c). As expected, ABE9 (evaluated 8954 sgRNAs) presented an extremely narrowed editing window of 1-2nt with the highest efficiency on position 5. Profiling of the motif preferences of the ABE9 showed that similar to ABE8e, they were suitable for a wide range of accurate A-to-G editing without strict motif requirements, suggesting their accuracy was dependent on the position relative to the protospacer but not on sequence context (Fig. 6d). As determined by thousands of sgRNAs, it suggests that ABE9 is very accurate to preferentially edit adenines in position 5 of the protospacers.

## Discussion

Highly efficient and precise correction of single nucleotide pathogenic mutation is demanded for gene therapy to reach its potential. Using structure-based design and molecular evolution of TadA-8e, we have generated ABE9, which efficiently edits adenines in a 1-2nt window without cytosine editing activity. To minimize the editing window of base editors, structure-based molecular evolution has been successfully leveraged to obtain new editors, such as BE4max-YE1 and YEE variants, which catalyze conversions within a 1-2 nucleotide window^35^, and eA3A-BE preferentially editing in a TCN motif^36^ and A3G-BEs selectively editing the second C in a CC motif^37^. Although ABEmax-F148A has been shown to reduce the editing window^7^, very limited effects have been observed when it has been transferred to TadA-8e (Fig. 1b), indicating the experiences from TadA7.10 could not be directly transferred to TadA-8e.

More complicated than CBEs, ABEs are capable of catalyzing both adenines and cytosines in a similar editing window^16^. Since the editing window of C-to-T is overlapped with that of A-to-G, it is impracticable to eliminate their cytosine deamination activity through reducing the editing window. While we were completing this project, Bae and colleagues reported introduction of D108Q in ABEmax or N108Q in ABE8e could reduce their cytosine deaminase activity^13^, which was consistent with our current study, suggesting the residue 108 was critical for the discrimination of substrates such as adenines and cytosines. The previous study also showed this residue was important for the recognition of ssDNA substrates as the D108N mutation was pivotal for the generation of eTadA, the unnatural DNA adenine deamination^3^. Moreover, the combinational mutation in ABEmax (TadA-E59A&N108W/Q) displayed greatly reduced RNA editing and preferentially catalyzing adenine conversions at protospacer position 5 but the activity was compromised^8^. It is consistent with our findings that ABE8e-N108Q exhibited reduced editing window and RNA off-target effects (Fig. 3b,d and Fig. 4c).

As for the discrimination between cytosines and adenines, we speculated that the mutation of N108 to a larger side chain residue (Q) would expel the backbone of its substrate. It apparently affected the deamination of cytosines greater than adenines since the pyrimidine ring of cytosines needs to be shifted further toward the pocket for the catalytic reaction to happen. However, TadA8e-N108Q still retained considerable cytosine deaminase activity (Fig. 3e,f) until the introduction of a second mutation, L145T, which fully abolished cytosine conversions and further narrowed the adenine editing window to 1-2nt without sacrifice on-target adenine conversion efficiency. We found that introduction of variant mutations at L145 had similar effects on reducing the editing window and cytosine bystander mutation as N108Q, suggesting the L145 position was a previously unnoticed residue which was also critical for substrate discrimination. The L145T mutation may adjust the pocket indirectly by influencing the positions of its nearby residues, P29, F84 and Y149, as L145 is located relatively far from the target base. F84 is also a critical residue identified in the initial generation of eTadA^3^. It is located within the pocket right below the target base ring and it forms a triangle platform together with Y149 and V28 to hold the base ring of the substrate. Additionally, we found that V28 could be a critical position involved in the discrimination of cytosines and adenines, since V28F and V28N showed a significant decrease of cytosine conversions, on the contrary, V28G had opposite effects (Fig. 1c), suggesting it is possible to innovate pure CBE, CGBE or dual-BE through further engineering of TadA-8e.

Developing a base editor with refined editing window is challenging, especially editing a specific base within promiscuous homopolymeric sites. Using selected target sites in cells and rodent embryos, we determined that ABE9 was accurate with a very narrow editing window. More importantly, through a guide-RNA target pair library containing over 9000 targets, the data showed that ABE9 could be considered as an ABE focusing on a 1-2nt editing window with the highest efficiency A_5_ (Fig. 6c). To our knowledge, it is potentially the most accurate ABE to date. As SpRY almost does not require any PAM sequence, ideally ABE9-SpRY could precisely target any adenine through an appropriate sgRNA for broad targeting scope. Importantly, ABE9 induces almost no off-target effects (either Cas9-dependent or -independent) at both DNA and RNA levels, which is an important feature not only for basic research but also critically important for clinical applications.

## Methods

### Plasmid construction

The primers and DNA sequences used in this research could be found in Supplementary Tables 1-3 and Supplementary Sequences. ABEmax (#112095) and ABE8e (#138489) were purchased from Addgene. PCR was performed using KOD-Plus-Neo DNA Polymerase (Toyobo, code no. KOD-401). Serial ABE plasmids generated in this article were constructed using ClonExpress MultiS One Step Cloning Kit (Vazyme). sgRNA expression plasmids were constructed as described previously^22, 38^. Briefly, oligonucleotides listed in Supplementary Table 1 were denatured at 95 °C for 5 min followed by slow cooling to room temperature (RT). Annealed oligonucleotides were ligated into BbsI-linearized U6-sgRNA(sp)-EF1α-GFP for sgRNA expression (Thermo Fisher Scientific).

### Human cell culture

HEK293T cell line (ATCC CRL-3216) was fostered in Dulbecco’s Modified Eagle’s medium (DMEM, Gibco), and DMEM medium was supplemented with 1% Penicillin-Streptomycin (Gibco) antibiotic mix and 10% (*v/v*) fetal bovine serum (FBS, Gibco). The cell line was maintained at 37 °C with 5% CO_2_ in the incubator.

### Cell transfection and genomic DNA extraction

HEK293T cells were seeded into 24-well plates (Corning) and cultured until the confluency was at approximately 80%. Next, 750 ng of ABEs and 250 ng of sgRNA expression plasmids were co-transfected using polyethyleneimine (PEI, Polysciences) following the manufacturer’s recommended protocol. Transfected cells were digested with 0.25% trypsin (Gibco) for sorting after three days and GFP+ cells were collected. And then cell genomic DNA was isolated using the TIANamp Genomic Kit (Tiangen Biotech) according to the manufacturer’s instructions. For rodent DNA extractions, mouse or rat tail tip genomic DNA was isolated using the One-Step Mouse Genotyping Kit (Vazyme) based on the manufacturer’s instructions.

### Modified orthogonal R-loop assay

Cas9-independent DNA off-target analysis in this study was using the modified orthogonal R-loop assay at each R-loop site with nSaCas9-sgRNA plasmid. For transfection, 250 ng of SpCas9 sgRNA plasmid, 300 ng of base editor plasmid (ABE8e, ABE8e-N108Q or ABE9) and 300 ng of nSaCas9-sgRNA plasmid were co-transfected into HEK293T cells using PEI. Transfected cells were digested with 0.25% trypsin (Gibco) for sorting after three days. Genomic DNA was isolated using the TIANamp Genomic Kit (Tiangen Biotech) according to the manufacturer’s instructions.

### Total mRNA preparation

For RNA off-target experiments, HEK293T cells were cultured in 10-cm dishes and transfected with 25 μg of Cas9n-P2A-GFP, ABE8e-P2A-GFP, ABE8e-N108Q-P2A-GFP or ABE9-P2A-GFP using PEI at approximately 80% confluency. Transfected cells were digested with 0.25% trypsin (Gibco) for sorting on FACSAria III (BD Biosciences) after three days. Cells were gated on their population via forward/sideward scatter after doublet exclusion (Supplementary Note). Then, ∼400,000 cells (top 15% GFP signal) were collected, and RNA was extracted according to standard protocols.

### RNA sequencing (RNA-Seq) experiments

A total of 3 μg RNA per sample was adopted as input material to get preparations for the sample. Sequencing libraries were generated using a NEBNext Ultra RNA Library Prep Kit for Illumina (NEB) following the manufacturer’s recommendations, and index codes were added to attribute sequences to each sample. Briefly, mRNA was purified from total RNA using poly-T oligo-attached magnetic beads. Fragmentation was carried out using divalent cations under elevated temperature in NEBNext First Strand Synthesis Reaction Buffer (5×). First-strand cDNA was synthesized using random hexamer primer and M-MuLV Reverse Transcriptase (RNase H-). Second-strand cDNA synthesis was subsequently performed using DNA Polymerase I and RNase H. Remaining overhangs were converted into blunt ends via exonuclease/polymerase activities. After adenylation of 3′ ends of DNA fragments, a NEBNext Adaptor with a hairpin loop structure was ligated to get the preparation for hybridization. In order to select cDNA fragments of preferentially 250-300bp in length, the library fragments were purified with the AMPure XP system (Beckman Coulter). 3 μl USER Enzyme (NEB) was then incubated with size-selected, adaptor-ligated cDNA at 37 °C for 15 min followed by 5 min at 95 °C before PCR. Next, Phusion High-Fidelity DNA polymerase, universal PCR primers and Index (X) Primer were used in PCR. Lastly, amplicons were purified (AMPure XP system) and library quality was assessed on the Agilent 2100 Bioanalyzer system. The clustering of index-coded samples was performed on a cBot Cluster Generation System using TruSeq PE Cluster Kit v3-cBot-HS (Illumina), according to the manufacturer’s instructions. The library preparations were sequenced on an Illumina HiSeq platform after cluster generation, and 125bp/150bp paired-end reads were obtained.

### Transcriptome-wide RNA analysis

For the RNA-Seq analysis, high sequencing data were firstly removed adapter sequences in reads with Trim Galore (version 0.6.6) (https://github.com/FelixKrueger/TrimGalore), and aligned with STAR^39^ (version 2.7.1a) to hg38 genome. Aligned BAMs were tag added, and sorted with SAMtools^40^ (version 1.14). Duplication removed with Picard MarkDuplicates module (version 2.23.9) (https://github.com/broadinstitute/picard) and filtered unmapped reads with SAMtools. Above BAMs were converted to mpileup format with SAMtools which records integrated mutation information. The significant mutation information was extracted based on mpileup files. The sites with coverage higher than 25, a mutation count at least 6 and mutation ratios over 5% were collected as filtered sites subsequently. As the edits found in mpileup files were filtered by removing the sites existing in the Cas9n-transfected condition, the sites only in Cas9n-transfected cases were the control.

### Animals and microinjection of zygotes

The manipulation in mouse embryos was performed as previously described^41^. To be brief, C57BL/6J, ICR strain mice and Sprague-Dawley strain rats purchased from the Shanghai Laboratory Animal Center were raised in standard cages in a special pathogen-free facility on a 12 h light/12 h dark cycle with a full supply of food and water. All animal experiments approved by the East China Normal University Center for Animal Research obeyed the regulations drafted by the Association for Assessment and Accreditation of Laboratory Animal Care in Shanghai. sgRNA with chemical modification was synthesized by GenScript (Nanjing, China), and mRNA was prepared as previously described^26^. The T7 promoter was introduced into ABE8e or ABE9 template by PCR using primers T7-mRNA (ABE8e/ABE9)-F/R (Supplementary Table 2). ABE8e or ABE9 mRNA was transcribed using the in vitro RNA transcription kit (mMESSAGE mMACHINE® T7 Ultra Kit, Ambion). mRNA was purified with MEGAclear™ Kit (Ambion). For microinjection, solutions containing complexes of ABE8e or ABE9 mRNA (100 ng μl^-1^) and sgRNA (200 ng μl^-1^) were diluted in nuclease-free water and injected into the cytoplasm by an Eppendorf TransferMan NK2 micromanipulator. Injected zygotes were immediately transferred into pseudopregnant female mice or rats after injection or after overnight culture in KSOM medium at 37 °C in a humidified atmosphere with 5% CO_2_.

### The creation of stable cell line disease models

The 150-bp fragments of G·C-to-A·T disease-associated genes from ClinVar database (https://www.ncbi.nlm.nih.gov/clinvar/) were assembled into a modified lenti-vector from lentiCRISPR v2 (#52961), obtaining transfer plasmids (Lenti *COL1A2*-EF1α-DsRed-P2A-puro, Lenti *CARD14*-EF1α-DsRed-P2A-puro, Lenti *BVES*-EF1α-DsRed-P2A-puro or Lenti *KCNA5*-EF1α-DsRed-P2A-puro). HEK293T cells were seeded into 24-well plates. After 12-16 h, cells were co-transfected with 300 ng transfer plasmids, 300 ng pMD2.G and 300 ng psPAX2 using PEI at approximately 80% confluency following the manufacturer’s instructions. Virus-containing supernatant was collected after 48 h of transfection, and then filtered through a 0.45-μm low protein binding membrane (Millipore). HEK293T cells were seeded into 12-well plates at approximately 40-50% confluency and 50 μl filtered virus-containing supernatant was added to the 12-well plates. After 24 h, cells transduced with lentivirus were split into new plate wells supplemented with puromycin (1 μg ml^-1^). After 72 h, cells were collected with the fewest existing colonies to obtain single-copy integration and then cultured for further transfection.

### The design of the library

The architecture of the oligos in the guide-target pair library was designed as previously described^33^. Each oligo contains a full-length sgRNA with a corresponding cassette targeted by the sgRNA. The spacers of the sgRNAs fulfilled the following criteria: (1) Each spacer is initiated with a Guanine; (2) The 4th to 9th positions of the spacers are composed of all possible 6-mers with at least an adenine or a cytosine. The 6-mers were surrounded by random 2-mers and 11-mers at 5’- and 3’-end, respectively. (3) Spacers with five consecutive thymines were avoided for it might impede the transcription. Each targeted cassette contained a 20-bp target sequence followed by an “NGG” PAM. Randomly selected wild-type human genomic sequences flanked the target sequence.

### Integration of the library and cell culture

The oligos were assembled into a modified pBlueScript backbone containing a U6 promoter and a hygromycin-resistant gene (hygro). The U6-sgRNA and hygro cassette were flanked by Tol2 sites to ensure its integration by Tol2 transposon. The construction and amplification of the library were finished by GENEWIZ Biotechnology Co. (Suzhou, China). For the library integration, HEK293T cells were seeded into 10-cm plates (Corning). We co-transfected the Tol2-transposon plasmid (10 μg) and library mixture (10 μg) into HEK293T cells at approximately 90% confluency. To facilitate sgRNA-integration, cells were selected with Hygromycin B (25 μg ml^-1^) (Thermo Fisher Scientific, CAT#10687010) one day after transfection, lasting for > 14 days, during which over 90% percent of cells were screened out. The screening was performed on at least 20 plates to ensure library coverage. When the HEK293T cells were at approximately 90% confluency over again, the second-round transfection was conducted by co-transfection of a Tol2-transposon plasmid (10 μg) and a base editor plasmid (ABE8e, ABE8e-N108Q or ABE9) (10 μg) that contains a blasticidin resistance gene and Tol2-transposase binding sites. The next day, 10 μg ml^-1^ Blasticidin S HCL (Thermo Fisher Scientific, CAT#A11139-03) was used for the second-round selection, lasting for >14 days. Again, the selection strength was adjusted so that over 90% of the cells were killed after 2-3 days of selection. When the density of cells was up to 80-90%, transfected cells were digested with 0.25% trypsin (Gibco) for genomic DNA extraction (see Library genomic DNA extraction). The target regions of libraries were amplified from genomic DNA (100 ng) by PCR with primers listed in Supplementary Table 2. The resulting amplicons were sequenced by the HTS platform of GENEWIZ Biotechnology Co. (Suzhou, China). The complete sequences of the mentioned plasmids and base editor sequences are appended in the Supplementary Sequences.

### Library genomic DNA extraction

Cells were digested with 0.25% trypsin and collected by centrifugation at 1,000 rpm for at least 3 min at RT. Cell pellets were resuspended and washed with proper volumes of PBS once followed by lysing with protein K at 55 °C for at least 1 h until the lysate became relatively clear. An equal volume of phenol-chloride was added into the lysate, followed by vortex for at least 1 min. The mixture was incubated at RT for at least 10 min for phase separation and was centrifuged at 12,000 rpm for at least 15 min. Carefully remove the aqueous layer into a new tube, add an equal volume of chloride, vortex for at least 1 min and incubate at RT for 10 min. The mixture was centrifuged again at 12,000 rpm for at least 15 min. Carefully remove the aqueous layer into a new tube again, add 1/10 volume of NaOAC (3.5 M) and 2.5 volume of ethanol, and incubate at -20 °C overnight. The following day samples were centrifuged at 12,000 rpm for at least 30 min at RT. Rinse the DNA pellet with 75% ethanol twice, and dissolve the DNA with Nuclease-Free Water (Ambion, AM9932).

### Editing efficiency calculation of the library and motif visualization

The JavaScript version of fastq-join (https://github.com/brwnj/fastq-join) firstly joins two fastq files from HTS. The combined fastq files were aligned to all of the amplicons in the library using BWA-mem (0.7.17-r1188) and the reads were divided for each amplicon to determine the connection between the amplicons and the sequenced reads. Reads with many equally plausible alignments were detected by the random mode. To minimize the influence of PCR amplification, targets with sequencing depths more than 20 times higher than the average depth of the library were abandoned for every library. Next, all of the reads to its corresponding amplicon pairwise were aligned by EMBOSS needle (https://www.ebi.ac.uk/Tools/psa/embossneedle/). Merely the reads that matched the following criteria were chosen for analysis: 10 bp sequences upstream and downstream of the 20 bp target sites completely matched the consensus sequences; the target sites included no detectable indels or degenerate base Ns. The editing type, the total number of reads aligned to amplicons, and the number of edited reads at each position were then analyzed to calculate each type’s absolute editing efficiency at each site. The relative editing efficiency was also computed relatively to the highest absolute editing efficiency. The matching sgRNA was accumulated once for each edited read when enriching motifs since the effectiveness of sgRNA varied greatly. The motifs edited at A_5_ were tallied and visualized using the “ggseqlogo” package in R.

### High-throughput DNA sequencing and data analysis

On- and off-target genomic regions of interest were amplified from genomic DNA (100-150 ng) using primers listed in Supplementary Tables 2 and 3. KOD-Plus-Neo DNA Polymerase and site-specific primers containing an adaptor sequence (forward 5′-GGAGTGAGTACGGTGTGC-3′; backward 5′-GAGTTGGATGCTGGATGG-3′) at the 5′ end were used for PCR to prepare high-throughput sequencing (HTS) libraries. The products mentioned were therefore subjected to a second-round PCR using primers with different barcode sequences. The amplified libraries were mixed and sequenced on an Illumina HiSeq platform. The A-to-G or C-to-T, A and G conversions and indels in the HTS data were analyzed by BE-Analyzer^23^.

### Statistics and reproducibility

Data are presented as mean ± s.d. from independent experiments. All statistical analyses were performed on at least n = 3 biologically independent experiments or three biologically independent samples unless otherwise noted in the figure captions. An unpaired two-tailed Student’s t-test was used to determine the significance of the differences between the two groups, using GraphPad Prism 9.3 (GraphPad Software). Specific *P* values are indicated in the figure captions. *P* < 0.05 was considered significant.

### Data availability

The outcome of HTS has been deposited in the NCBI Sequence Read Archive database under accession codes PRJNA812697 and PRJNA812700. RNA-seq data have been deposited in the NCBI Sequence Read Archive database under accession code PRJNA811343. Data for rat embryos treated with ABE7.10 have already been posted in the NCBI Sequence Read Archive database under accession code PRJNA471163 from the previous study. Source data for Figs. 1-6 and Extended Data Figs. 1-7 is presented with the paper. All other data supporting the discoveries of this research are available from the corresponding author on reasonable request.

## Acknowledgments

We are grateful to Stefan Swiko (Texas A&M University Health Science Center) for proof reading of the manuscript and the East China Normal University Public Platform for innovation (011). We thank Ying Zhang from the Flow Cytometry Core Facility of School of Life Sciences in ECNU and Haixia Jiang from the Core Facility and Technical Service Center for SLSB of School of Life Sciences and Biotechnology in SJTU. We thank Li Ji (MedSci) for drawing schematic diagrams. This work was partially supported by grants from National Key R&D Program of China (2019YFA0110802 and 2019YFA0802800), the National Natural Science Foundation of China (No.32025023, No.31971366), and grants from the Shanghai Municipal Commission for Science and Technology (21CJ1402200, 20140900200), the Innovation Program of Shanghai Municipal Education Commission (2019-01-07-00-05-E00054), the Fundamental Research Funds for the Central Universities, and the support from East China Normal University Outstanding Doctoral Students Academic Innovation Ability Improvement Project (YBNLTS2021-026).

## Author contributions

L.C. and D.L. designed the experiments. L.C., S.Z., N.X., M.H., X.Z., J.Y., S.B., Y.H., C.L., B.Z., G.R., M.L. and L.W. performed the experiments. L.C., S.Z., N.X., M.H., X.Z., J.Y., S.B., Y.H., H.M., H.W., D.Z., C.Y., M.L., Y.C. and D.L. analyzed the data, D.L., L.C., S.Z., N.X., Z.X., D.Z. and G.S. wrote the manuscript with the input from all the authors, D.L. supervised the research.

## Figure legends

**Extended Data Fig. 1.**
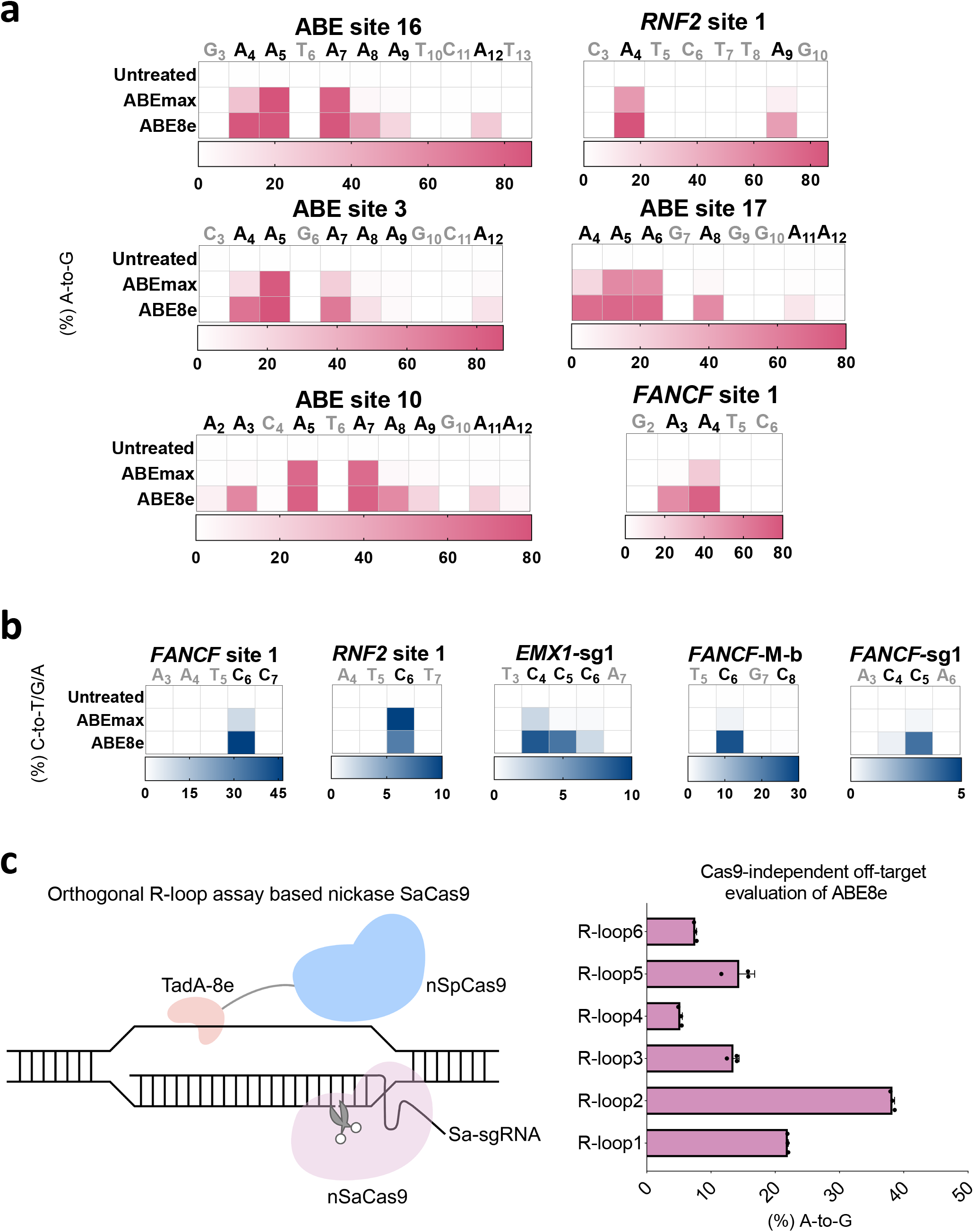
ABE8e induces severe bystander mutations and global random off-target editing. **a**, Comparison ng efficiency of ABEmax or ABE8e at 6 endogenous genomic loci in HEK293T cells. Partial data are derived, c. **b**, The C-to-T/G/A editing efficiency of ABEmax or ABE8e was examined at 5 endogenous genomic loci in ells. Partial data are derived from Fig. 1c. **c**, The schematic diagram of orthogonal R-loop assay-based nickase Cas9) (left panel); Cas9-independent DNA off-target analysis of ABE8e using the modified orthogonal R-loop R-loop site with nSaCas9-sgRNA plasmid (right panel). Data are mean ± s.d. (n = 3 independent experiments). ived from Fig. 4b. In **a** and **b**, the heatmap represents average editing percentage derived from three independent and editing efficiency was determined by HTS. Statistical source data are provided online.

**Extended Data Fig. 2.**
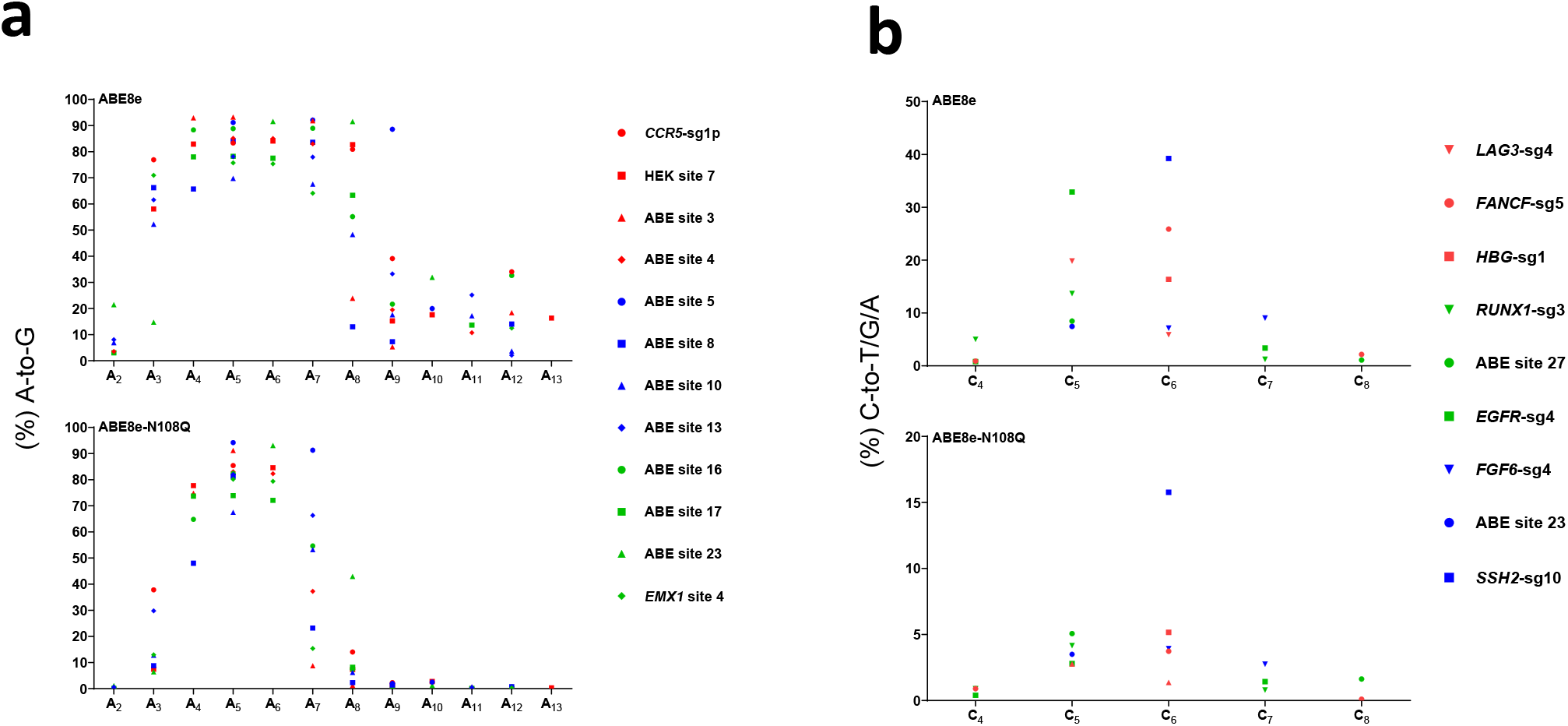
Comparison of the editing window between ABE8e and ABE8e-N108Q. **a**, Comparison of A-to- g window of ABE8e or ABE8e-N108Q at 12 target sites in HEK293T cells. **b**, Comparison of C-to-T/G/A base w of ABE8e or ABE8e-N108Q at 9 target sites in HEK293T cells. In **a** and **b**, data are from Fig. 2a (**a**) and nd each point represents means from three independent experiments. Statistical source data are provided online.

**Extended Data Fig. 3.**
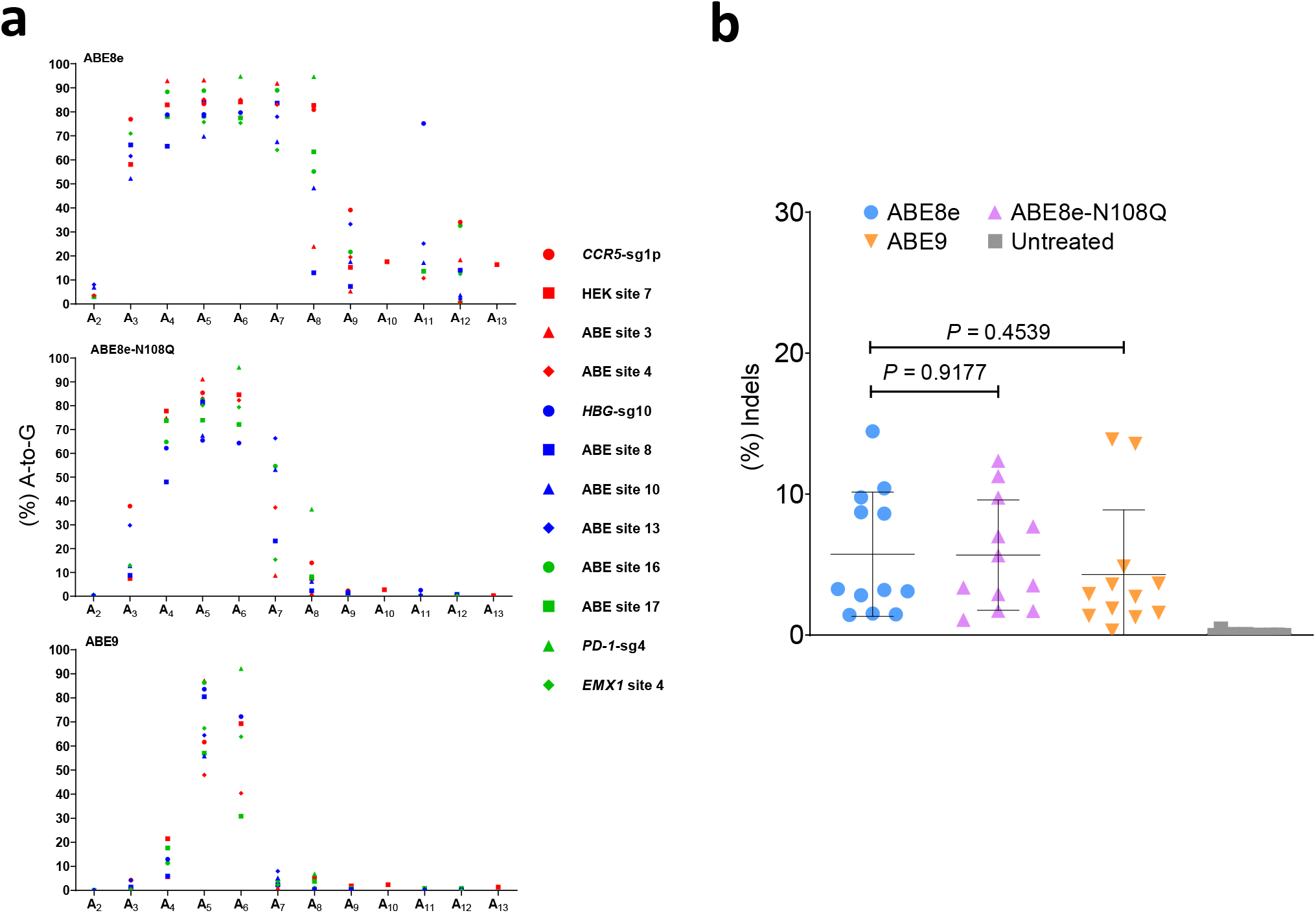
Characterization of A-to-G base editors of ABE8e, ABE8e-N108Q and ABE9. **a**, Comparison of editing window of ABE8e, ABE8e-N108Q or ABE9 at 12 target sites in HEK293T cells. Data points represent three independent experiments. Data are derived from Fig. 3b. **b**, Comparison of indels induced by ABE8e, 8Q or ABE9 at 12 target sites from Fig. 3b. Each data point represents the average indel frequency at each target d from 3 independent experiments. Error bar and *P* value are derived from these 12 data points. *P* value was y two-tailed Student’s t test. Statistical source data are provided online.

**Extended Data Fig. 4.**
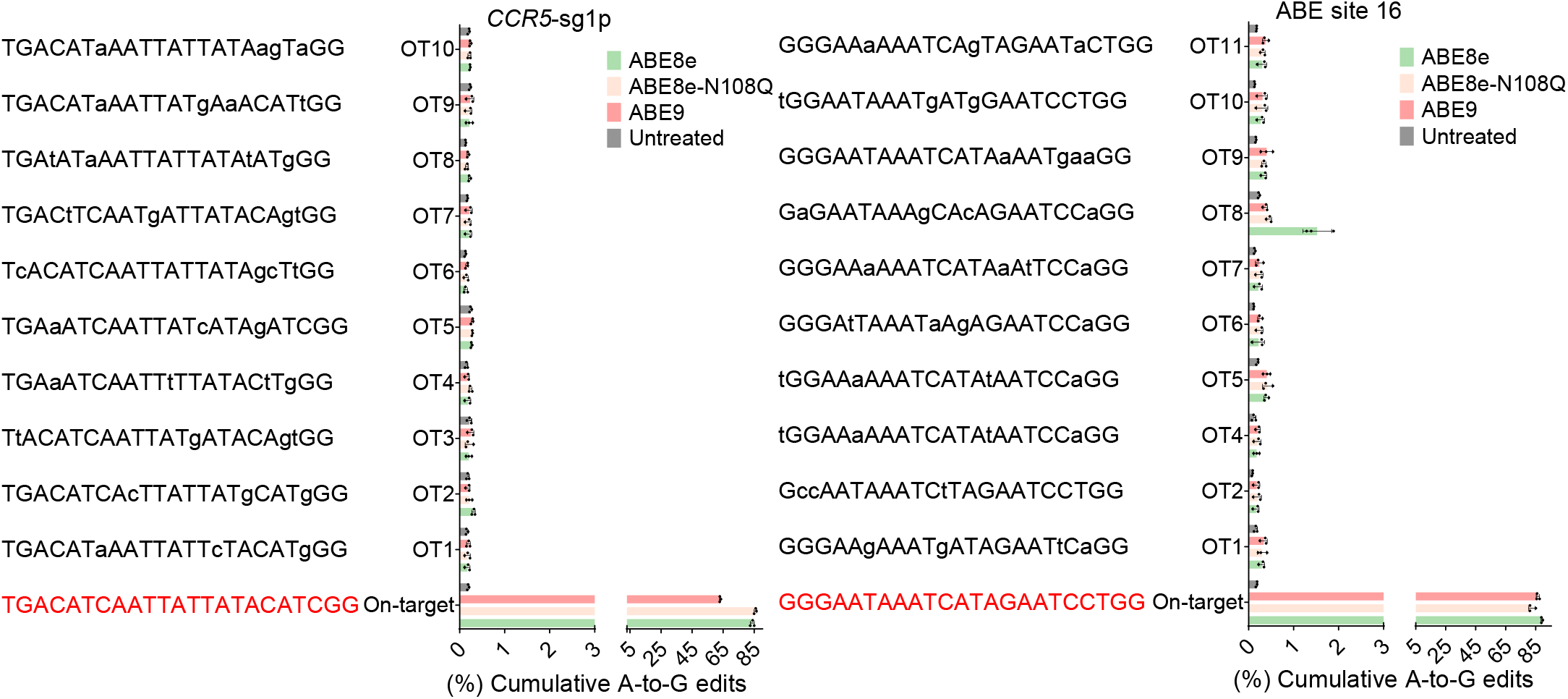
Cas9-dependent off-target assessment of ABE9. Cas9-dependent DNA on- and off-target analysis of targets (*CCR5-*sg1p and ABE site 16) by ABE8e, ABE8e-N108Q and ABE9 in HEK293T cells. Data are = 3 independent experiments). On-target data are derived from Fig. 3b. Statistical source data are provided online.

**Extended Data Fig. 5.**
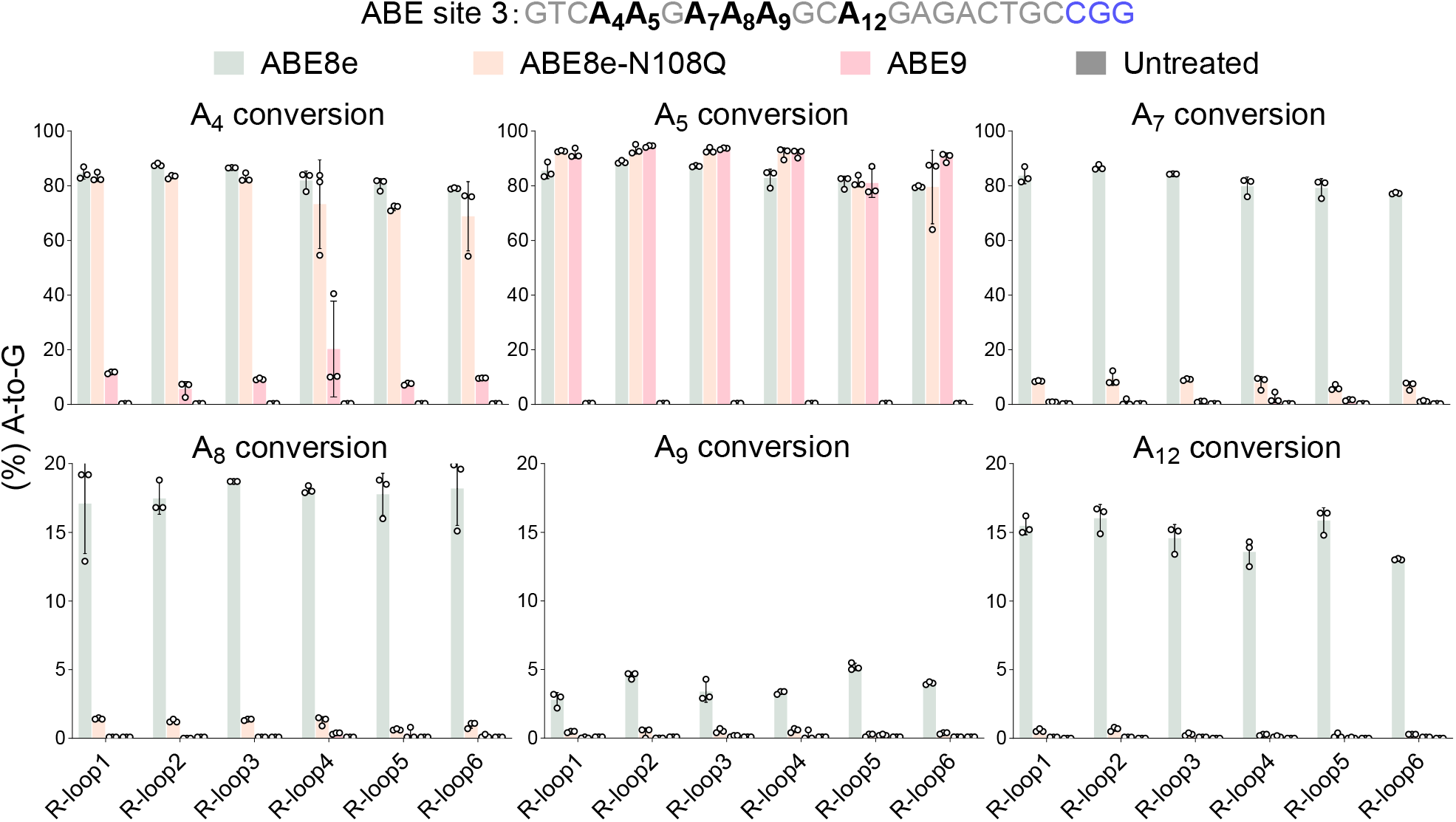
On-target base editing by ABE9 in the R-loop assay. On-target base editing induced by ABE8e, 8Q or ABE9 using the modified orthogonal R-loop assay at each R-loop site with nSaCas9-sgRNA plasmid. an ± s.d. (n = 3 independent experiments). Statistical source data are provided online.

**Extended Data Fig. 6.**
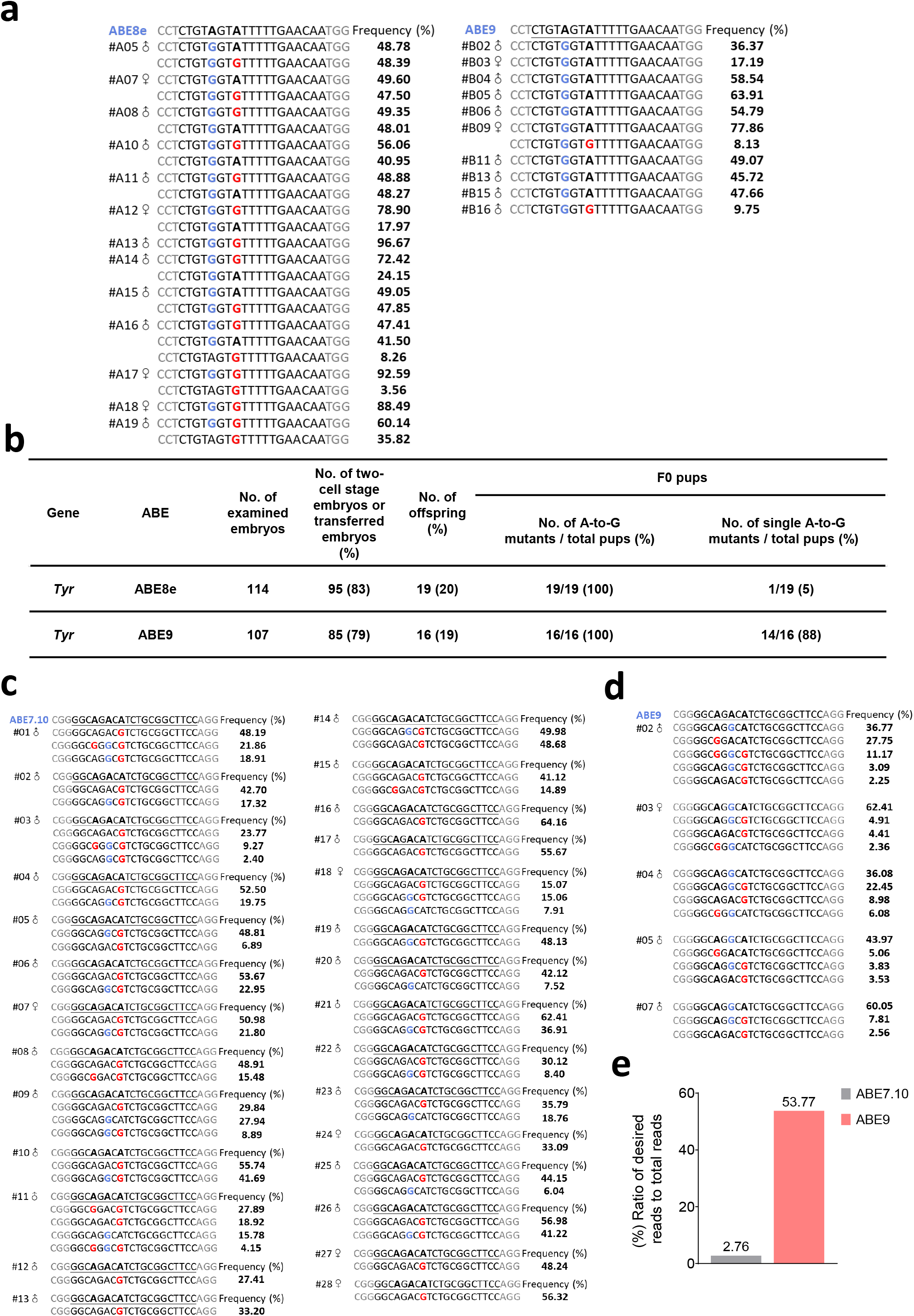
Highly efficient and precise editing by ABE9 in rodent embryos. **a**, Genotyping of F0 generation y ABE8e or ABE9. **b**, Summary of the numbers of embryos used and the pups generated after microinjection of A or ABE9/sgRNA. **c, d**, Genotyping of F0 rats induced by ABE7.10 (**c**) and ABE9 (**d**) (desired editing in blue diting in red). **e**, Ratio of desired reads to total reads in F0 rats induced by ABE7.10 or ABE9. In **a, c** and **d**, the the right represents the frequency of the indicated mutant allele. Percentiles of each allele reads <1% are omitted. ce data are provided online.

**Extended Data Fig. 7.**
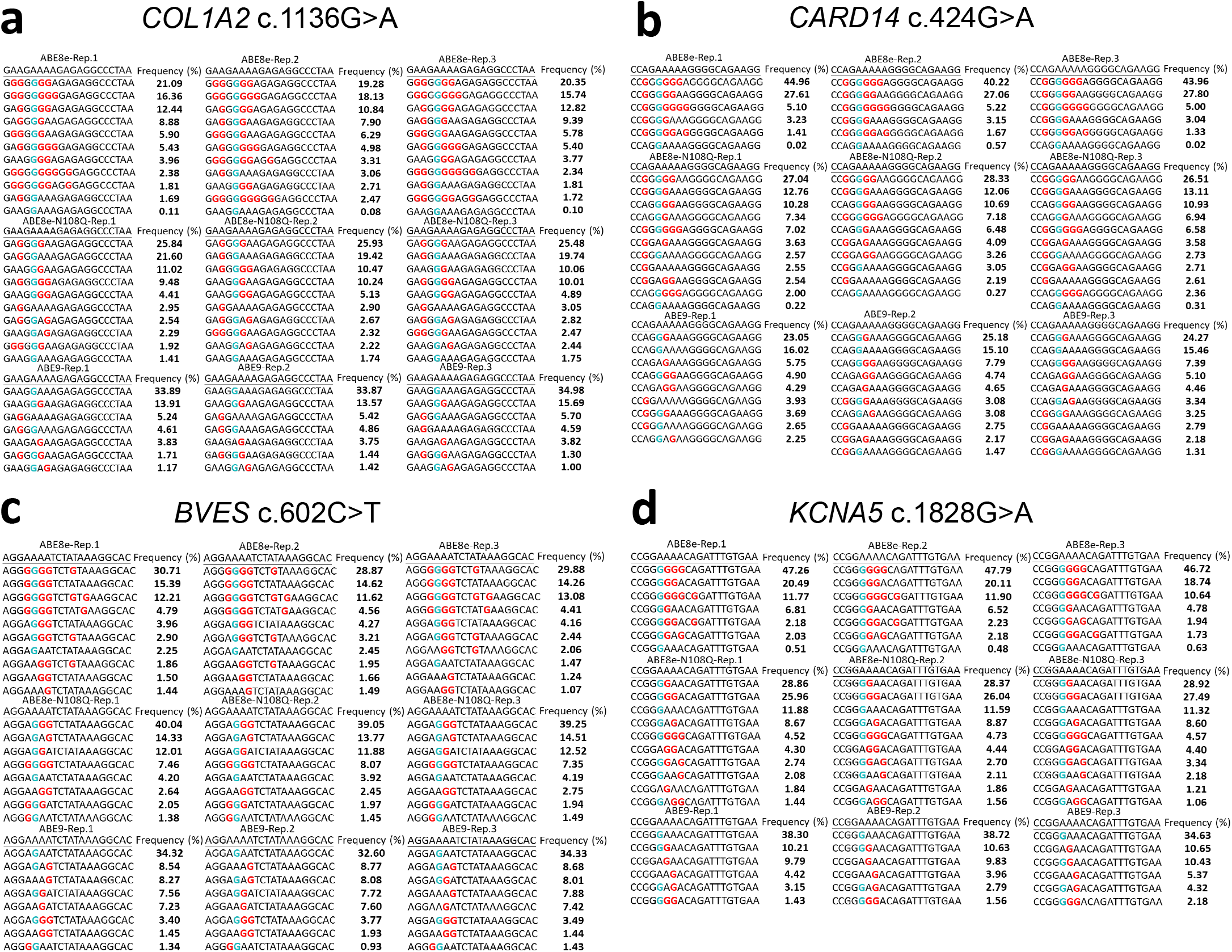
Allele tables for ABE9 in four stable HEK293T cell lines. **a-d**, Allele tables for ABE8e, ABE8e-BE9 in four stable HEK293T cell lines: *COL1A2* c.1136G>A (**a**), *CARD14* c.424G>A (**b**), *BVES* c.602C>T (**c**) 1828G>A (**d**). The percentile and sequencing reads of each allele from two or three independent experiments are ight. Desired A_5_-to-G percentiles of alleles are exhibited, while percentiles of top ten invalid allele types are percentiles of invalid allele types less than 1% are omitted.

## References

1. Rees, H.A. & Liu, D.R. Base editing: precision chemistry on the genome and transcriptome of living cells. Nat. Rev. Genet. 19, 770–788 (2018).

2. Komor, A.C., Kim, Y.B., Packer, M.S., Zuris, J.A. & Liu, D.R. Programmable editing of a target base in genomic DNA without double-stranded DNA cleavage. Nature 533, 420–424 (2016).

3. Gaudelli, N.M. et al. Programmable base editing of A*T to G*C in genomic DNA without DNA cleavage. Nature 551, 464–471 (2017).

4. Zuo, E. et al. Cytosine base editor generates substantial off-target single-nucleotide variants in mouse embryos. Science 364, 289–292 (2019).

5. Jin, S. et al. Cytosine, but not adenine, base editors induce genome-wide off-target mutations in rice. Science 364, 292–295 (2019).

6. Grunewald, J. et al. Transcriptome-wide off-target RNA editing induced by CRISPR-guided DNA base editors. Nature 569, 433–437 (2019).

7. Zhou, C. et al. Off-target RNA mutation induced by DNA base editing and its elimination by mutagenesis. Nature 571, 275–278 (2019).

8. Rees, H.A., Wilson, C., Doman, J.L. & Liu, D.R. Analysis and minimization of cellular RNA editing by DNA adenine base editors. Sci. Adv. 5, eaax5717 (2019).

9. Richter, M.F. et al. Phage-assisted evolution of an adenine base editor with improved Cas domain compatibility and activity. Nat. Biotechnol. 38, 883–891 (2020).

10. Gaudelli, N.M. et al. Directed evolution of adenine base editors with increased activity and therapeutic application. Nat. Biotechnol. 38, 892–900 (2020).

11. Musunuru, K. et al. In vivo CRISPR base editing of PCSK9 durably lowers cholesterol in primates. Nature 593, 429–434 (2021).

12. Newby, G.A. et al. Base editing of haematopoietic stem cells rescues sickle cell disease in mice. Nature 595, 295–302 (2021).

13. Jeong, Y.K. et al. Adenine base editor engineering reduces editing of bystander cytosines. Nat. Biotechnol. 39, 1426–1433 (2021).

14. Li, S., Liu, L., Sun, W., Zhou, X. & Zhou, H. A large-scale genome and transcriptome sequencing analysis reveals the mutation landscapes induced by high-activity adenine base editors in plants. Genome Biol. 23, 51 (2022).

15. Xu, L. et al. Efficient precise in vivo base editing in adult dystrophic mice. Nat. Commun. 12, 3719 (2021).

16. Kim, H.S., Jeong, Y.K., Hur, J.K., Kim, J.S. & Bae, S. Adenine base editors catalyze cytosine conversions in human cells. Nat. Biotechnol. 37, 1145–1148 (2019).

17. Lee, H.K. et al. Targeting fidelity of adenine and cytosine base editors in mouse embryos. Nat. Commun. 9, 4804 (2018).

18. Liu, Z. et al. Efficient generation of mouse models of human diseases via ABE- and BE-mediated base editing. Nat. Commun. 9, 2338 (2018).

19. Lapinaite, A. et al. DNA capture by a CRISPR-Cas9-guided adenine base editor. Science 369, 566–571 (2020).

20. Doman, J.L., Raguram, A., Newby, G.A. & Liu, D.R. Evaluation and minimization of Cas9-independent off-target DNA editing by cytosine base editors. Nat. Biotechnol. 38, 620–628 (2020).

21. Wang, L. et al. Eliminating base-editor-induced genome-wide and transcriptome-wide off-target mutations. Nat. Cell Biol. 23, 552–563 (2021).

22. Zhang, X. et al. Increasing the efficiency and targeting range of cytidine base editors through fusion of a single-stranded DNA-binding protein domain. Nat. Cell Biol. 22, 740–750 (2020).

23. Hwang, G.H. et al. Web-based design and analysis tools for CRISPR base editing. BMC Bioinformatics 19, 542 (2018).

24. Kim, K. et al. Highly efficient RNA-guided base editing in mouse embryos. Nat. Biotechnol. 35, 435–437 (2017).

25. Ryu, S.M. et al. Adenine base editing in mouse embryos and an adult mouse model of Duchenne muscular dystrophy. Nat. Biotechnol. 36, 536–539 (2018).

26. Yang, L. et al. Increasing targeting scope of adenosine base editors in mouse and rat embryos through fusion of TadA deaminase with Cas9 variants. Protein Cell 9, 814–819 (2018).

27. Landrum, M.J. et al. ClinVar: public archive of interpretations of clinically relevant variants. Nucleic Acids Res. 44, D862–868 (2016).

28. Symoens, S. et al. Type I procollagen C-propeptide defects: study of genotype-phenotype correlation and predictive role of crystal structure. Hum. Mutat. 35, 1330–1341 (2014).

29. Forlino, A., Cabral, W.A., Barnes, A.M. & Marini, J.C. New perspectives on osteogenesis imperfecta. Nat. Rev. Endocrinol. 7, 540–557 (2011).

30. Jordan, C.T. et al. Rare and common variants in CARD14, encoding an epidermal regulator of NF-kappaB, in psoriasis. Am. J. Hum. Genet. 90, 796–808 (2012).

31. Schindler, R.F. et al. POPDC1(S201F) causes muscular dystrophy and arrhythmia by affecting protein trafficking. J. Clin. Invest. 126, 239–253 (2016).

32. Yang, Y. et al. Novel KCNA5 loss-of-function mutations responsible for atrial fibrillation. J. Hum. Genet. 54, 277–283 (2009).

33. Arbab, M. et al. Determinants of Base Editing Outcomes from Target Library Analysis and Machine Learning. Cell 182, 463–480 e430 (2020).

34. Wang, D. et al. Optimized CRISPR guide RNA design for two high-fidelity Cas9 variants by deep learning. Nat. Commun. 10, 4284 (2019).

35. Kim, Y.B. et al. Increasing the genome-targeting scope and precision of base editing with engineered Cas9-cytidine deaminase fusions. Nat. Biotechnol. 35, 371–376 (2017).

36. Gehrke, J.M. et al. An APOBEC3A-Cas9 base editor with minimized bystander and off-target activities. Nat. Biotechnol. 36, 977–982 (2018).

37. Lee, S. et al. Single C-to-T substitution using engineered APOBEC3G-nCas9 base editors with minimum genome- and transcriptome-wide off-target effects. Sci. Adv. 6, eaba1773 (2020).

## References

38. Zhang, X. et al. Dual base editor catalyzes both cytosine and adenine base conversions in human cells. Nat. Biotechnol. 38, 856–860 (2020).

39. Dobin, A. et al. STAR: ultrafast universal RNA-seq aligner. Bioinformatics 29, 15–21 (2013).

40. Li, H. et al. The Sequence Alignment/Map format and SAMtools. Bioinformatics 25, 2078–2079 (2009).

41. Li, D. et al. Heritable gene targeting in the mouse and rat using a CRISPR-Cas system. Nat. Biotechnol. 31, 681–683 (2013).

